# Repeatability analysis improves the reliability of behavioral data

**DOI:** 10.1101/2019.12.19.882043

**Authors:** Juliane Rudeck, Silvia Vogl, Stefanie Banneke, Gilbert Schönfelder, Lars Lewejohann

## Abstract

Reliability of data has become a major concern in the course of the reproducibility crisis. Especially when studying animal behavior, confounding factors such as novel test apparatus can lead to a wide variability of data. At worst, effects of novelty or stress related behavior can mask treatment effects and the behavioral data may be misinterpreted. Habituation to the test situation is a common practice to circumvent novelty induced increases in variance and to improve the reliability of the respective measurements. However, there is a lack of published empirical knowledge regarding reasonable habituation procedures and a method validation seems to be overdue.

This study aimed at setting up a simple strategy to increase reliability of behavioral data. Therefore, exemplary data from mice tested in an Open Field (OF) arena were used to elucidate the potential of habituation and how reliability of measures can be confirmed by means of a repeatability analysis using the software R. On seven consecutive days, male C57BL/6J, BALB/cJ and 129S1/SvImJ mice were tested in an OF arena once daily and individual mouse behavior (distance travelled, average activity) was recorded. A repeatability analysis was conducted in order to estimate the reliability of measured animal behavior with regard to repeated trials of habituation.

Our data analysis revealed that monitoring animal behavior during habituation is important to determine when individual differences of the measurements are stable. Here, the mixed effect model framework proved to be a powerful tool for estimating repeatability values. Repeatability values from distance travelled and average activity increased over the habituation period, revealing that around 60 % of the variance of the data can be explained by individual differences between mice. The first day of habituation was statistically significantly different from the following 6 days in terms of distance travelled and average activity. A habituation period of three days appeared to be sufficient in this study. Overall these results emphasize the importance of habituation and in depth analysis of habituation data to define the correct starting point of the experiment for improving the reliability and reproducibility of experimental data.

## 1 Introduction

Animal experimentation always requires an ethical evaluation, balancing the scientific benefit and possible constraints to animal welfare. On the one hand, the generation of high-quality research data is crucial for a better translation of experimental data to humans. On the other hand, stress and distress in laboratory animals should be limited to a minimum. Fortunately, these two goals are often working perfectly together because it is well known that data derived from non-stressed animals is of higher quality expressed, for example, in the reduction of the standard deviation and/ or a higher consistency of scientific data (Bell et al., 2009; Gouveia and Hurst, 2017).

One well known possibility to minimize stress and distress for the laboratory animal is the prior habituation to unknown experimental equipment or handling procedures (Gouveia and Hurst, 2017; Toval et al., 2017). The term habituation originates from behavioral biology and describes the diminution of a response induced by constant or repeated exposure to a novel stimulus and displays a simple form of non-associative learning (Leussis and Bolivar, 2006). A distinction is made between habituation over time (intrasession) and over repeated exposures (intersession) (Leussis and Bolivar, 2006). In this original field of behavioral biology, the habituation process is the primary research objective and serves as a model for non-associative learning and memory mechanisms as well as for pharmacological studies (Leussis and Bolivar, 2006; Pavkovic et al., 2018; Evans et al., 2019). One of the most frequently assessed parameters of habituation is a decrease in exploratory behavior measured in the Open Field (OF) test as a change in distance travelled or activity. Once the animal is familiar with the new environment, the explorative behavior is reduced and it is considered that the laboratory animal has habituated and the learning and memory process is completed (Leussis and Bolivar, 2006).

Knowledge about habituation can be used to introduce a wide variety of behavioral tests and to increase the intended performance in the actual test situation (Bowen et al., 2016; Gouveia and Hurst, 2017; Toval et al., 2017; Moreton et al., 2019). For example, in rodents, physical activity can be measured under voluntary or forced condition. On the one hand, voluntary exercise has the disadvantage of very variable volume and intensity differing from animal to animal. On the other hand, rodents in forced running wheel tests show low level of performance. Habituation and training with the running wheel have been shown to improve locomotor performance in the forced running wheel system in rats and consequently increased the validity of the scientific data (Toval et al., 2017). Although, such effects of habituation are frequently applied, the documentation in the literature is heterogeneous. In case of the OF, there are research groups which use this test without documented habituation (e.g., (Singh et al., 2015; Wirz et al., 2015; Breu et al., 2016; Sichova et al., 2018; Teixeira et al., 2018). Some of them stated that they want to test for novelty induced activity (Breu et al., 2016). Other researchers establish only one day of habituation to either reduce novelty induced activity or to measure the effect of habituation from trial one to trial two (Simoes et al., 2017; de Sousa et al., 2018; de Melo et al., 2019). Frequently, experimenters place the animals 30 min prior to testing in the corresponding room or arena and define this as habituation (Pagani et al., 2015; Nikolaus et al., 2016; Aso-Someya et al., 2018). Rarely, a previous habituation is mentioned in the literature which is carried out for more than two days (Milanovic et al., 2016; Nasehi et al., 2017; Chellammal et al., 2019). However, there is usually no well-founded derivation of the habituation period which is stated in the literature. A strategy how to determine an effective habituation period for a certain experiment would be favorable.

The repeatability analysis is one possible method to assess the accuracy of measurements which can be normally or not normally distributed (Nakagawa and Schielzeth, 2010) and it allows identifying an explanation for the occurring variances. Repeatability describes the proportion of the total variation that is reproducible among the repeated measurements of the same group (Shrout and Fleiss, 1979; Lessells and Boag, 1987; Nakagawa and Schielzeth, 2010). Nakagawa and Schielzeth published a practical guide for biologists to bring forward the method of repeatability analysis in the scientific community (Nakagawa and Schielzeth, 2010). This method of data analysis is commonly employed in the field of ecology and evolutionary biology where the repeatability of morphological, physiological and behavioral traits is of special interest (Araya-Ajoy et al., 2015). Some examples include the wing morphology of drosophila (Houle et al., 2003), cancer evolution (Taylor et al., 2013), or mate choice for reproduction (Pogany et al., 2014; Haneke-Reinders et al., 2017; Burley et al., 2018). More recently, the concept of animal personality that focuses on consistent between-individual differences in behavior, was investigated by applying repeatability analysis (Koski, 2011; Dingemanse and Dochtermann, 2013; Brent et al., 2014; Brust et al., 2015; Roche et al., 2016; Chen et al., 2018). In this context, the repeatability between several trails on one day or successive days as well as between several independent experiments of different research groups is meant. Bell et al. showed in a meta-analysis of repeatability of independent experiments from different research groups for animal behavior that many behavioral types are more consistent within individuals than previously assumed (Bell et al., 2009). In addition, Mazzamuto et al. underline the importance of method validation to ensure that the intended parameter is really captured (Mazzamuto et al., 2019). Given that there is growing concern regarding reproducibility of experimental data in biomedical science (Baker, 2016), we are convinced that a documented validation of the habituation data is a beneficial approach in all fields of animal behavioral science to increase the reliability and reproducibility of experimental data. A mathematically justified and comprehensible explanation for the choice of a certain habituation period would strengthen the scientific outcome, as it is shown that the intended parameter can actually be captured. In addition, valuable information can be obtained from the habituation data e.g., the source of variance, which might be helpful for the interpretation of the primary experimental results.

Therefore, the aim of our study was to elucidate the putative hidden information of habituation data in combination with the repeatability analysis. As an example, we investigated retrospectively the effectiveness of intersession habituation of male mice of C57BL/6J, BALB/cJ and 129S1/SvImJ inbred strains to an OF arena within two different experiments carried out at the same institute (one in winter and one in summer time). For this purpose we analyzed video data to assess locomotor parameters in the OF (distance travelled, average activity). We conducted a repeatability analysis to check the reliability of our data from trial to trial and to evaluate if the chosen habituation period was sufficient or could be reduced in future studies. In order to identify the primary source of variance in locomotion data, we examined the factors experiment/ batch, strain, and the individual animal (animal ID). All these factors are known from the literature to affect locomotion (Podhorna and Brown, 2002; Leppanen et al., 2006; Bell et al., 2009).

## 2 Material and Methods

### 2.1 Ethics statement

The data for this study is based on two experiments which were approved by the Berlin state authority, *Landesamt für Gesundheit und Soziales*, under license No. G 0309/15 and G 0194/16 and were conducted in accordance with the German Animal Protection Law (TierSchG, TierSchVersV).

### 2.2 Animals and animal care

Male C57BL/6J mice (n = 38) from Charles River (Sulzfeld, Germany, breeding the original Jackson strain from the USA) and male BALB/cJ mice and male 129S1/SvImJ mice (n = 15 per strain) from Jackson Laboratory (Bar Harbor, USA, imported via Charles River) were obtained after weaning, to familiarize them with the housing conditions. The mice were free of all viral, bacterial, and parasitic pathogens listed in the Federation of European Laboratory Animal Science Associations (FELASA) guidelines. During familiarization time, mice were housed in groups of 5 - 8 animals per cage in Eurostandard type III polycarbonate cages with filter tops (Tecniplast, Hohenpeissenberg, Germany), autoclaved bedding and nesting material (LASbedding™PG2, LASvendi, Soest, Germany), a mouse house consisting of cardboard (LBS Biotechnology, United Kingdom) and gnawing material as environmental enrichment (J. Rettenmaier & Söhne GmbH + Co KG, Rosenberg, Germany). The animals had free access to autoclaved food pellets (LASQCdiet™ Rod16, LASvendi, Soest, Germany) and water acidified with HCL (pH 2.5 – 3.0 to prevent growth of algae and pathogens). The room temperature was maintained at 21 ± 1 °C, with a relative humidity of 55 ± 10 %. The light/ dark cycle in the room consisted of 12/ 12 h artificial light with lights on from 5.00 am to 5.00 pm in winter time and 6.00 am to 6.00 pm in summer time. All animals were handled by cupping the mouse in open hands.

### 2.3 Habituation

After two weeks of familiarization to the housing conditions, all mice were single housed to prevent aggressive behavior. With the start of the habituation period, mice were six to seven weeks old. On seven consecutive days, each mouse was habituated once a day to the following experimental conditions in a randomized order. Each mouse was fixated by grip in the neck to mimic the stress of a subcutaneous (s.c.) injection and was subsequently placed in a round OF arena (ø 30 cm, 40 cm height, 109 lux) for 5 min, monitored by video camera from above. Afterwards, the mouse was placed on the warm (at 35 °C) Incremental Hot Plate (IHP, IITC Inc. Life Science, Woodland Hills, USA) for 4 min, monitored by two video cameras (frontal and lateral). Habituation and testing took place between 9.00 am to 11.00 am in winter time and 10.00 am to noon in summer time to avoid influence by circadian rhythm. On day one and six, body weight of each mouse was measured within the routine animal check as a general welfare indicator (supplementary material Figure S1). Since there are hints in literature that the sex of the experimenter might have an influence (Bohlen et al., 2014), the whole habituation and experiments were performed by the same female experimenter. The results from pain assessment on the IHP will not be part of this publication.

### 2.3 Data collection and analysis

Locomotor activity, monitored as distance travelled (cm), and average activity (%, threshold 0.1 cm/s)were assessed for every mouse for 3 min in the round OF arena and analyzed with the Viewer Software (version 4, Biobserve GmbH, Bonn, Germany). We have chosen the mean of the last three minutes of observation for the evaluation to exclude possible initial stress behavior.

Statistical analysis and plotting of the graphs were conducted with the freely available software R, version 3.5.1 and R-studio, version 1.1.383 (www.r-project.org) and with the software Graph Pad Prism (version 6, Graph Pad Software, San Diego, USA). To get an overview of the data set, all data points of distance travelled and average activity were plotted with R and statistically analyzed with Graph Pad Prism using the Friedman test with Dunn’s multiple comparison test compared to day one. To test the repeatability of the data and to identify the source of the variance within the data, a repeatability analysis was performed using the “rptR”library to quantify the constancy of phenotypes (Nakagawa and Schielzeth, 2010; Stoeffel et al., 2018). First, data sets were tested for normal distribution by Q-Q-norm-plot (supplementary material Figure S2). If data did not show substantial deviation from normal distribution the repeatability value R was calculated using a linear mixed-effect model (LMM) based on Gaussian distribution. Animal ID, strain or experiment/ batch were included as random factors, whereas, distance travelled (cm) or average activity (%) served as fixed factors. The confidence interval (CI) [2.5 %, 97.5 %] is based on 500 bootstrapping runs and 100 permutations. To test for statistical significance of the repeatability of distance travelled or average activity between the selected numbers of trials, the likelihood ratio test was performed. To identify the most important random factor, R was first calculated over the seven-day habituation period for all fixed factors including animal ID, strain and experiment as multiple random grouping factors. To observe the progress of repeatability over the whole time, R was calculated additionally over three adjacent measurements (days), including the random grouping factors animal ID or strain. In a second step, the identified effect covariate strain that explain only a part of the variances in the data was included to adjust the repeatability for the predominating random factor animal ID.

## 3 Results

### 3.1. Effects of habituation and buprenorphine on OF behavior of single mice

In the initial step, the raw data of each animal were examined.

From the second day of habituation onwards, a considerable reduction of distance travelled and average activity occurred compared to day one (Figure 1). In addition, the widths of confidence intervals of distance travelled and average activity decreased from day one until reaching stable plateaus at day five (Table 1).

**Figure 1:**
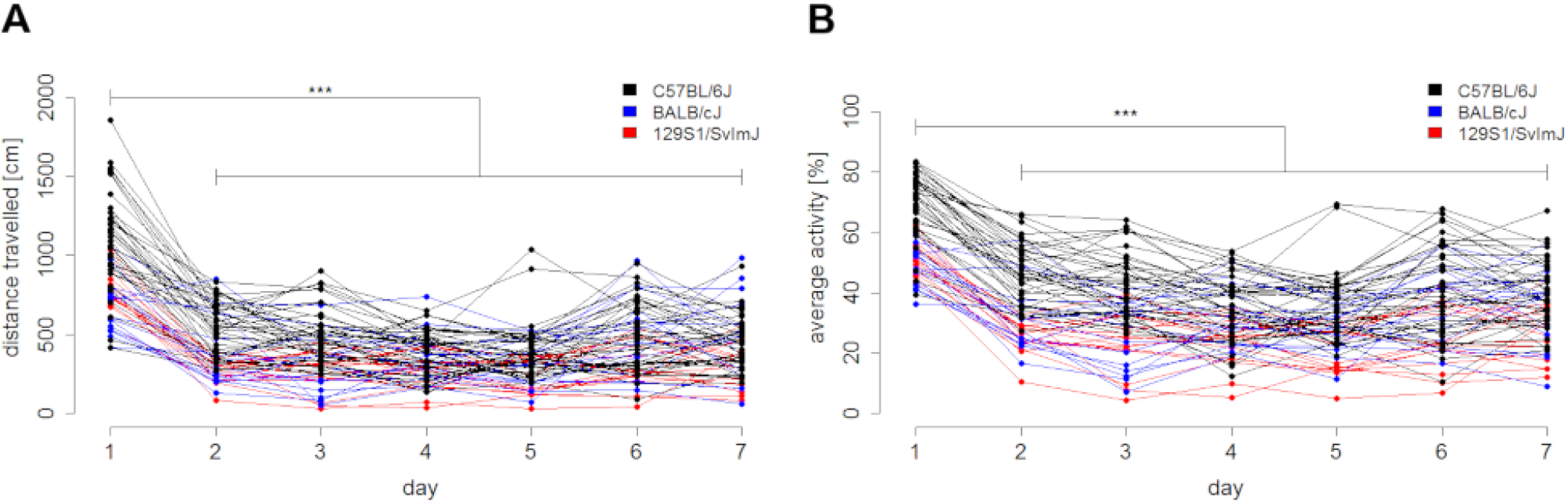
Frequency of observed behavior decreases during habituation. (A) Distance travelled [cm] and (B) average activity [%] are depicted. Values are presented as exact value per each animal (n = 38 C57BL/6J, n = 15 BALB/cJ and n = 15 129S1/SvImJ male mice). The Friedman test with Dunn’s multiple comparisons test was used for statistical evaluation (n.s. = not significant (A) p-value = 0.7119, (B) p-value > 0.9999, **** p-value < 0.0001 (A, B).

**Table 1:** Variances of distance travelled and average activity within the habituation period. The [2.5 %. 97.5 %] confidence interval was calculated for the factor travelled distance and average activity for ever day of the habituation period (n = 38 C57BL/6J, n = 15 BALB/cJ and n = 15 129S1/SvImJ male mice).

|  | day 1 | day 2 | day 3 | day 4 | day 5 | day 6 | day 7 |
| --- | --- | --- | --- | --- | --- | --- | --- |
| distance travelled [cm] | [863.5, 1016] | [402.2, 497.1] | [353.9, 449.3] | [317.1, 387.8] | [297.9, 379.3] | [389.6, 497.9] | [372.7, 468.3] |
| average activity [%] | [58.09, 64.65] | [35.68, 42.03] | [31.33, 38.04] | [29.15, 34.54] | [28.04, 33.66] | [32.81, 40.12] | [33.08, 39.02] |

### 3.2. Repeatability analysis of explorative behavior

In a second step, a repeatability analysis was carried out in order to obtain reliable evidence for the successful habituation and additional information on the variance of the data.

In order to identify the main factor by which the observed variance in locomotion data can be explained, a repeatability analysis with multiple grouping factors was conducted over the seven-day habituation period (Table 2). The distance travelled as well as the average activity served as fixed factors and animal ID, strain, and experiment/ batch functioned as random multiple grouping factors. The variances observed for distance travelled and average activity is primarily explained by animal ID and strain. The fact that we included data from two different experiments did not contribute significantly to the observed variances of distance travelled or average activity (Table 2).

**Table 2:** Repeatability values with multiple grouping factors: animal ID, strain and experiment. For every factor, the repeatability R, the [2.5 %, 97.5 %] confidence intervals (CI) and the p-values, calculated by likelihood ratio test, were displayed over the seven-day habituation period (n = 38 C57BL/6J, n = 15 BALB/cJ and n = 15 129S1/SvImJ male mice). Estimation of repeatability was conducted with a linear mixed-effect model. The CI resulted from 500 bootstrapping runs and 100 permutations.

|  | repeatability for animal ID |  |  | repeatability for strain |  |  | repeatability for experiment |  |  |
| --- | --- | --- | --- | --- | --- | --- | --- | --- | --- |
|  | R | CI | p-value | R | CI | p-value | R | CI | p-value |
| distance travelled | 0.101 | [0.034, 0.184] | 1.31E-4 | 0.1 | [0, 0.298] | 0.00219 | 0.0124 | [0, 0.094] | 0.296 |
| average activity | 0.137 | [0.065, 0.241] | 3.6E-08 | 0.173 | [0, 0.443] | 7.59E-05 | 0.053 | [0, 0.232] | 0.0606 |

For this reason, repeatability values for distance travelled and average activity were calculated including both, animal ID and strain as a random factor. To record changes in repeatability over the whole period of habituation, each repeatability value R was calculated over three adjacent days resulting in five groupings (Table S1, S2). Within the first grouping (day 1 – 3) repeatability values of 0.027 (distance travelled, random factor: ID) and 0.177 (distance travelled, random factor: strain) as well as 0.199 (activity, random factor: ID) and 0.321 (activity, random factor: strain) was calculated for distance travelled and average activity, respectively (Figure 2). This indicates that only 2.7 % of the variance of distance travelled data and 19.9 % of the variance of the average activity data can be explained by the factor animal ID. 17.7 % of the variance of distance travelled data and 32.1 % of the average activity data can be explained by the factor strain. The repeatability of the results within this period is considered unlikely. In addition, most of the variance cannot be explained by definable factors and is therefore considered random noise likely induced by other factors, e.g. stress or anxiety-related behavior. From the second (day 2 – 4) to the fifth grouping (day 5 – 7) stable repeatability values of 0.556, 0.617, 0.505, and 0.498 for distance travelled (random factor: ID) and 0.634, 0.652, 0.627, and 0.604 for average activity (random factor: ID) were calculated (Figure 2 A, B). This means that roughly 50 to 60 % of the variance can be explained by individual differences between the mice (Table S1, S2). In contrast, only 27.6, 22.9, 18.2, 19.0 % and 40.1, 36.3, 29.7, 30.7 % of the observed variance in distance travelled and average activity data, respectively, can be traced back to strain (Figure 2 C, D, Table S1, S2).

**Figure 2:**
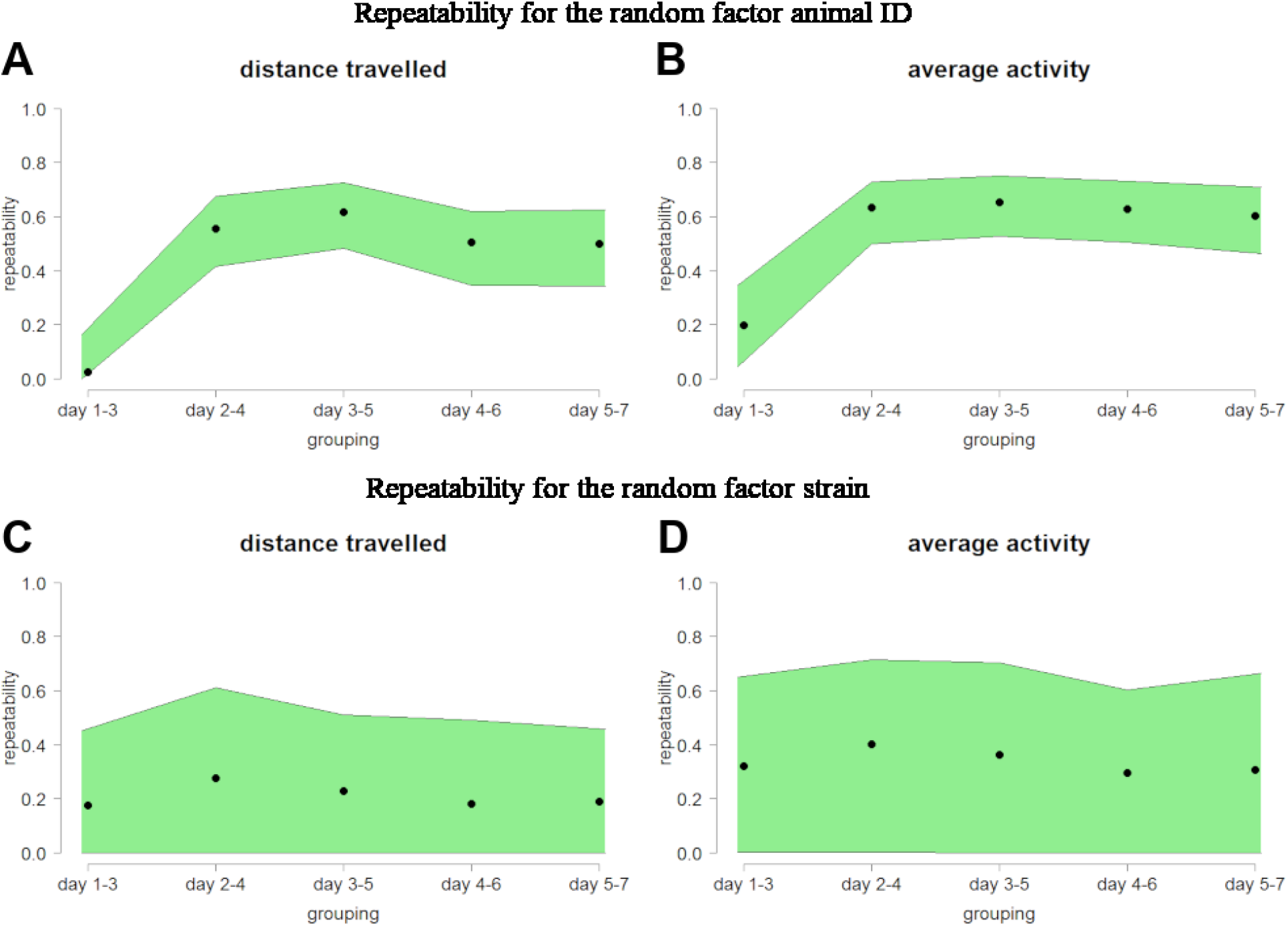
Animal ID and strain repeatability increase during habituation. (A, B) Calculated animal ID repeatability value and (C, D) strain repeatability value for the factors (A, C) distance travelled and (B, D) average activity were presented. Each repeatability value (R, black points) was calculated over three adjacent days resulting in five groupings (n = 38 C57BL/6J, n = 15 BALB/cJ and n = 15 129S1/SvImJ male mice). Estimation of repeatability was conducted with a linear mixed-effect model. The [2.5 %, 97.5 %] confidence intervals (CI) were displayed in green, resulting from 500 bootstrapping runs and 100 permutations.

From this repeatability analysis it can be concluded that strain is an important factor, but the influence of the individually stable behavior of each mouse predominates. Hence, estimations of repeatability over the whole time were calculated again including only animal ID as random factor with adjustment for strain (Figure 3, Table S1). The resulting adjusted repeatability values are comparable to animal ID repeatability values for distance travelled and average activity showing only slightly reduced (~15 %) repeatability values.

**Figure 3:**
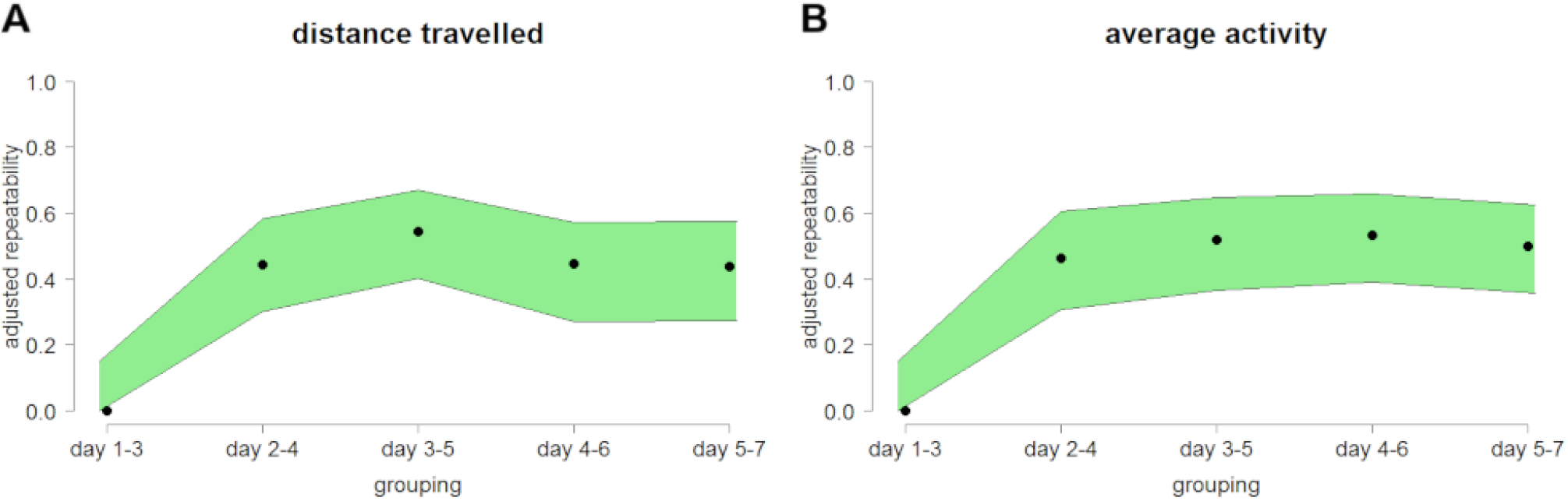
Animal ID repeatability adjusted for strain increase during habituation. Calculated animal ID repeatability values adjusted for the fixed factor strain were presented for (A) distance travelled and (B) average activity. Each repeatability value (R, black points) was calculated over three adjacent days resulting in five groupings (n = 38 C57BL/6J, n = 15 BALB/cJ and n = 15 129S1/SvImJ male mice). Estimation of repeatability was conducted with a linear mixed-effect model for locomotion as well as activity and with a generalized linear mixed-effect model for rearing and sniffing behavior. The [2.5 %, 97.5 %] confidence intervals (CI) were displayed in green, resulting from 500 bootstrapping runs and 100 permutations.

## 4 Discussion

Measured data of each individual mouse as well as adjusted repeatability values showed a stable pattern after three days of habituation for distance travelled and average activity. Moreover, additional habituation days did not lead to a further reduction of explorative behavior or increased repeatability of investigated parameters and reached a plateau in the second grouping from day 2 to day 4. Hence, a habituation time of three days seemed to be sufficient for this experimental set up. Furthermore, the individual personality of the mouse has a greater influence on the variance of the data than the strain to which it belongs. On the basis of these results a simple strategy can be established to introduce habituation as a preliminary step in behavioral experiments (Figure 4). First, plan and set up the experiment and habituate the animals to the testing situation. The simultaneous or retrospective checking of the measured data with repeatability analysis allows the determination of the optimal starting point of the experiment.

**Figure 4:**
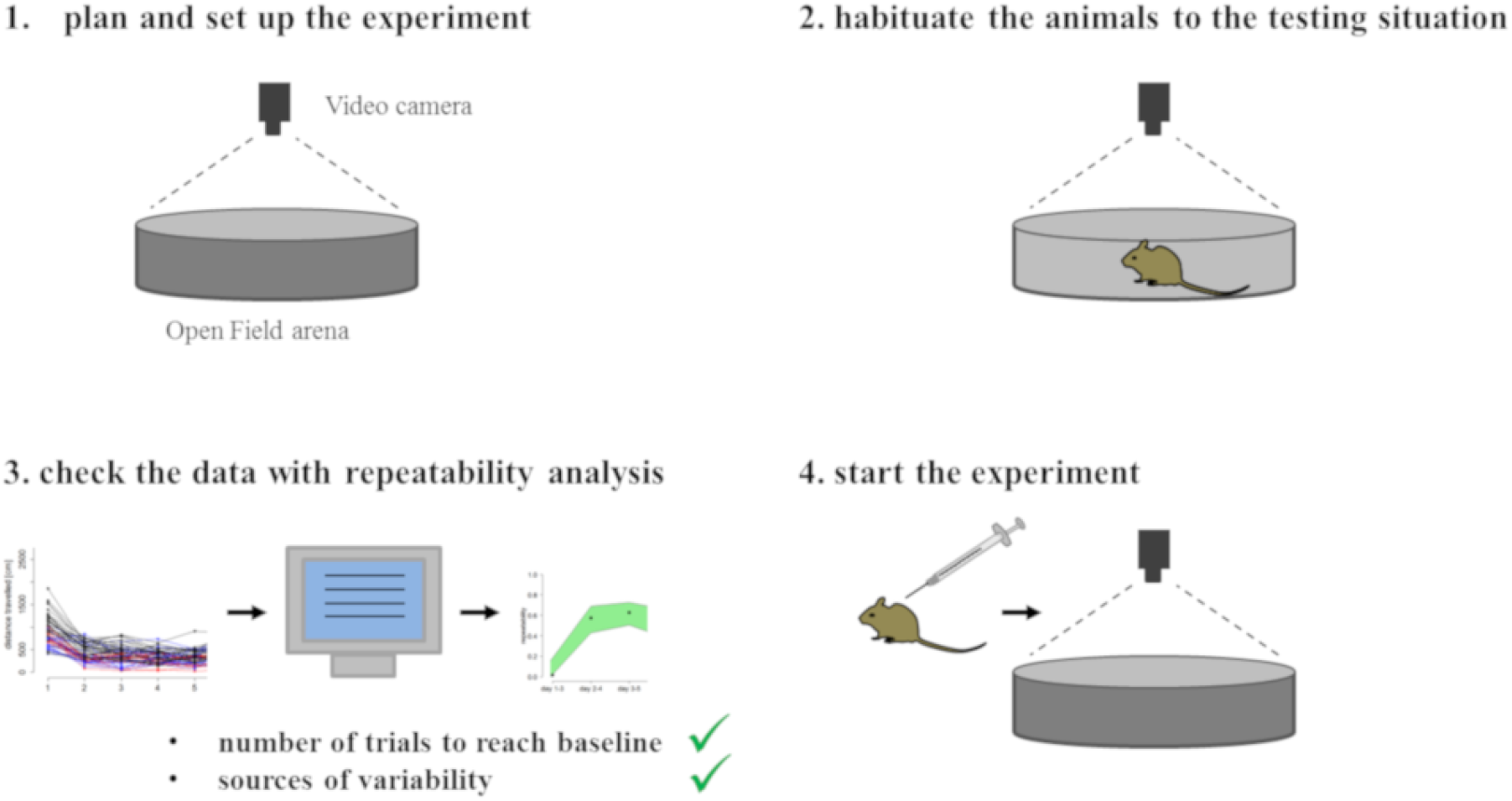
Strategy for the implementation of behavioral experiments. This basic strategy can be applied to all experiments with laboratory animals to determine the optimal starting point of your experiment.

In this study, a direct measurement of stress-related behavior or stress hormones was not performed. As reduction in body weight can give a hint of the general well-being of the animal, we measured the body weight on day one and six of habituation as part of the routine check-up. No loss of body weights was detected and, hence, we conclude that multiple habituation days to the least did not influence this general welfare parameter. Comparable to this result, Bodden et al. investigated the impact of repeated versus single open-field testing on welfare in C57BL/6J mice and also found no body weight changes or other signs of compromised welfare (Bodden et al., 2018). The anxiety of an animal is usually connected to time spent in the center and time spent nearby the wall of an OF (Leussis and Bolivar, 2006). We have not assessed these parameters because of the small size of the OF used and for this reason cannot comment on the anxiety behavior of the animals. In our study, we were able to demonstrate that the habituation approach was successful in decreasing the distance travelled and average activity.

Furthermore, we could show that repeatability analysis is a powerful tool to assess reproducible animal behavior and to identify the source of variance within the measured data (Nakagawa and Schielzeth, 2010). Even if the variance of the data cannot be reduced completely, knowing the sources of variability allows a much better interpretation of behavioral data. In connection with the reproducibility crisis, Baker reported that more than half of the researchers participating in a Nature’s survey have failed to reproduce their own experiments (Baker, 2016). The two experiments used here as examples were carried out with an interval of nine month. Our analysis did not reveal any significant effect of time of experimentation, indicating that our approach yield reproducible results. Even if the variance of locomotion data can be only reduced to a certain extent, the variance becomes explainable from the second day. Based on the results of the repeatability analysis, the influence of data from two different experiments could be excluded as the source of variation within this data set. For this reason we were able to pool the data and concentrated on the other two scientifically relevant factors, namely strain and the individual animal.

Interestingly, within the first grouping the variance of the data could not be traced back to either strain or the individual animal and is therefore more likely induced by unknown other factors, e.g., stress or anxiety. Only after the second grouping reliable parameters have been identified by the repeatability analysis as the source of variance. From this we concluded that, at the minimum, one day, better three days of habituation would be advisable for this or comparable set ups to be able to observe reproducible behavior. Dealing with reproducible behavior, on the other hand, supports a reliable interpretation of the variance of the data with regard to contribution of strain or individual animal personality.

Mouse strain comparisons are commonly implemented in behavioral and genetic studies with widely use of C57BL/6 and BALB/c mice (Crawley and Davis, 1982). However, previous reports on explorative behavior of C57BL/6J and BALB/c mice are inconsistent (Crawley and Davis, 1982; Belzung and Griebel, 2001; Augustsson and Meyerson, 2004; Brown et al., 2011). C57BL/6J mice were found to be engaged in more exploratory activity than BALB/cJ mice (Augustsson and Meyerson, 2004; Brown et al., 2011). In contrast, other studies reported higher anxiety levels associated with lower activity in C57BL/6 than BALB/c mice (Avgustinovich et al., 2000). An and colleagues reported no differences in locomotor activity between male C57BL/6J and BALB/cJ mice in a novel cage observation test (An et al., 2011). In our data set only a small percentage of the variance in explorative data could be traced back to strain. From this we concluded that strain differences were not the primary source of variance using these three strains. The large confidence interval might be explained by the fact that only three different strains have been used.

Another possible explanation is that there is a wide variation in the individual personality of mice, regardless of the strain. In recent years, behavioral studies focused more on consistent between-individual differences giving evidence that the animal personality in addition to strain is a crucial factor to explain variance of measurements (Koski, 2011; Lewejohann et al., 2011; Brent et al., 2014; Brust et al., 2015; Roche et al., 2016; Chen et al., 2018). This assumption is supported by a metaanalysis for animal behavior of Bell and colleagues. They demonstrated that many behavioral types are more consistent within individuals than previously assumed (Bell et al., 2009). Interestingly, such individual differences are also found in genetically homogeneous inbred mice housed under highly standardized conditions. This indicates that individual differences tend to escape standardization approaches and might reflect an evolutionary intrinsic value that perpetuates inter-individual differentiation (Lewejohann et al., 2011). In addition, Brust and colleagues showed that the personality of a mouse stabilizes with age and is highly repeatable over their lifespan (Brust et al., 2015). In our study, we observed highly consistent between-individual differences for distance travelled and average activity from the second grouping onwards. Interestingly, the individual differences between the mice, i.e., their “animal personality” (Lewejohann et al., 2011) was identified as the main contributor to variability in explorative behavior in the repeatedly conducted OF test. However, if only a single OF test was conducted, the variability in the data could not have been fully explained as the results of the OF test were only reasonably repeatable after habituation. Consequently testing without habituation might have had a considerable effect on the recently found lack of reproducibility, known as the reproducibility crisis. In accordance to these findings, Chappell and colleagues observed variable locomotion behavior but highly repeatable between-individual differences in mice (Chappell et al., 2004). Integrating the concept of animal personality not only offers the possibility to improve the interpretation and thus the quality of scientific data, but also to enhance animal welfare science by solving welfare problems on an individual level (Richter and Hintze, 2019).

### 4.1 Conclusion

We demonstrated a simple strategy to improve the data quality of behavioral experiments by implementing the repeatability analysis method to prove successful habituation as well as explaining underlying factors for variability in the data. While the overall variability is not necessarily reduced, detailed analysis of habituation data reveals the sources of variability. Furthermore, analyzing the habituation data by means of a repeatability analysis helps to determine the optimal start point of an experiment. In our data, the variances observed in the behavior patterns were predominantly due to the individual personality of the mice rather than a strain dependent effect. Behavioral data was repeatable after successful habituation and calculated repeatability values reached a plateau after only a few habituation trials. Hence, as a refinement strategy, the habituation period can be reduced to three days in future experiments of the same design. Taken together, this exemplary study underlines the importance of publishing and evaluating habituation data for all kind of behavioral experiments. Importantly, declaration of the effectiveness of habituation procedures could furthermore enhance the validity of experimental data and fortify reproducibility.

## 5 Acknowledgments

This work was supported by the Federal Ministry of Education and Research (BMBF) ‘[grant number 031A262D]’ and by the DFG [FOR 2591; LE 2356/5-1].

## 6 Authors contributions statement

JR, LL conceptualized the research. JR performed the experiments, conducted data analysis, wrote the main manuscript text and prepared the figures. GS wrote the BMBF proposal and raised the BMBF funding. LL wrote the DFG proposal and raised the DFG funding. JR, SV, SB, GS, LL revised the manuscript.

