## Supplemental material for "Repeatability analysis improves the reliability of behavioral data"

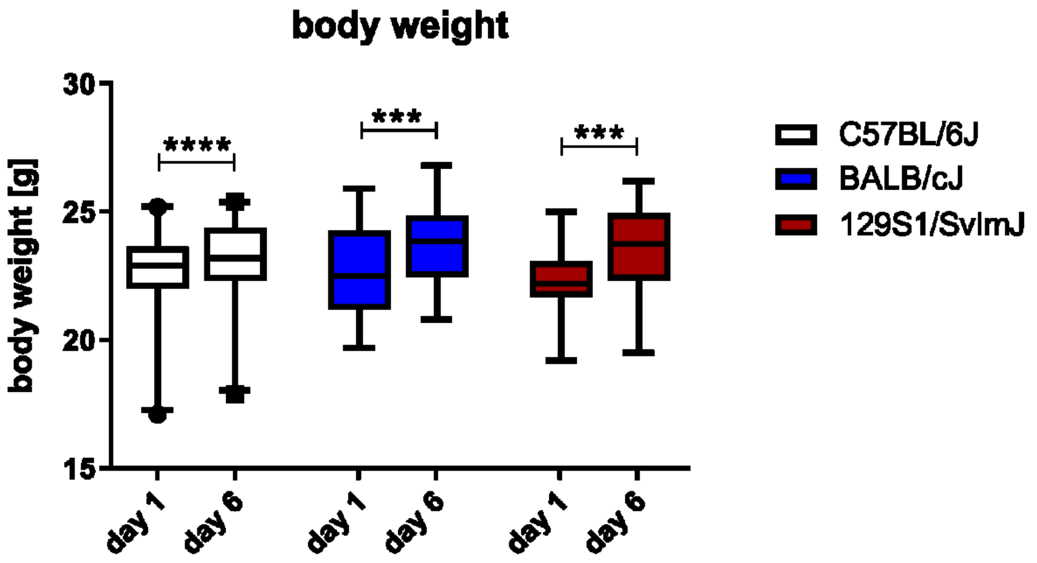


**Figure S1: Slightly increasing body weight during habituation period**
Body weight [g] at day one and six of habituation period per strain were presented as box plot with median and whiskers [2.5, 97.5 %] (n = 38 C57BL/6J: p-value = 0.0018, n = 15 BALB/cJ: p-value = 0.0004 and n = 15 129S1/SvImJ: p-value < 0.0001, Wilcoxon matched-pairs signed rank test, two-tailed).

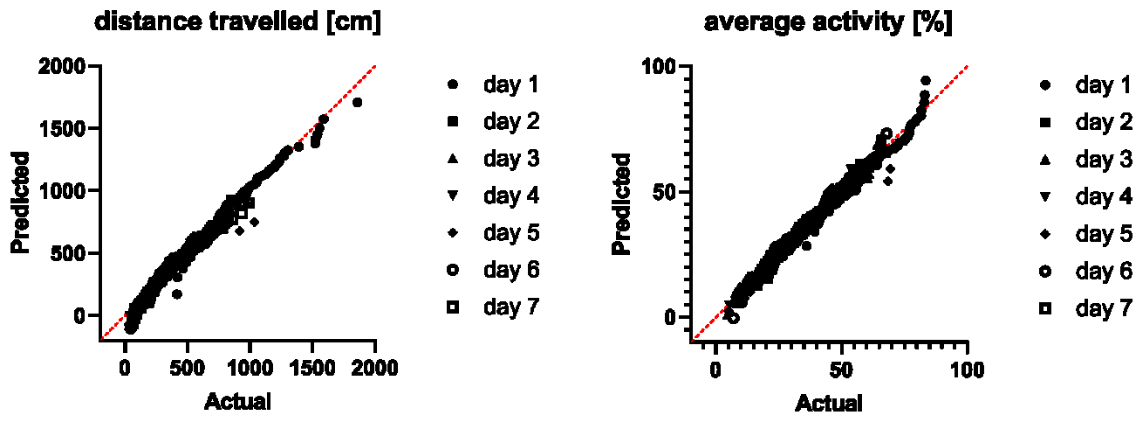


**Figure S2: Testing for normal distribution**
Data sets of distance travelled and average activity were checked for normal distribution using Q-Q-norm plot (n = 38 C57BL/6J, n = 15 BALB/cJ and n = 15 129S1/SvImJ male mice).

**Table S1: Repeatability values for animal ID as random factor with and without adjustment for the factor strain.**
Each repeatability value (R) was calculated over three adjacent days resulting in five groupings. For every factor R, the [2.5 %, 97.5 %] confidence intervals (CI) and p-values calculated by likelihood ratio test were displayed (n = 38 C57BL/6J, n = 15 BALB/cJ and n = 15 129S1/SvImJ male mice). Estimation of repeatability was conducted with a linear mixed-effect model. The CI resulted from 500 bootstrapping runs and 100 permutations.

| | **distance travelled** | | | **adjusted distance travelled** | | | **average activity** | | | **adjusted average activity** | | |
|---|---|---|---|---|---|---|---|---|---|---|---|---|
| **grouping** | **R** | **CI** | **p** | **R** | **CI** | **p** | **R** | **CI** | **p** | **R** | **CI** | **p** |
| **day 1-3** | 0.027 | [0, 0.166] | 0.377 | 0 | [0, 0.15] | 0.5 | 0.199 | [0.046, 0.345] | 0.00413 | 0 | [0, 0.15] | 1 |
| **day 2-4** | 0.556 | [0.415, 0.675] | 9.99E-15 | 0.445 | [0.302, 0.584] | 2.66E-09 | 0.634 | [0.501, 0.729] | 2.17E-19 | 0.465 | [0.307, 0.605] | 4.52E-10 |
| **day 3-5** | 0.617 | [0.482, 0.726] | 2.79E-18 | 0.544 | [0.403, 0.669] | 1.3E-13 | 0.652 | [0.528, 0.75] | 1.08E-20 | 0.519 | [0.365, 0.649] | 2.05E-12 |
| **day 4-6** | 0.505 | [0.347, 0.62] | 3.16E-12 | 0.445 | [0.271, 0.572] | 2.52E-09 | 0.627 | [0.506; 0.730] | 6.53E-19 | 0.534 | [0.391, 0.657] | 4.37E-13 |
| **day 5-7** | 0.498 | [0.344, 0.626] | 6.49E-12 | 0.438 | [0.274, 0.575] | 4.58E-09 | 0.604 | [0.465, 0.710] | 1.94E-17 | 0.499 | [0.357, 0.626] | 1.7E-11 |

**Table S2: Repeatability values for strain as random factor**
Each repeatability value (R) was calculated over three adjacent days resulting in five groupings. For every factor R, the [2.5 %, 97.5 %] confidence intervals (CI) and p-values calculated by likelihood ratio test were displayed (n = 38 C57BL/6J, n = 15 BALB/cJ and n = 15 129S1/SvImJ male mice). Estimation of repeatability was conducted with a linear mixed-effect model. The CI resulted from 500 bootstrapping runs and 100 permutations.

| | **distance travelled** | | | **average activity** | | |
|---|---|---|---|---|---|---|
| **grouping** | **R** | **CI** | **p** | **R** | **CI** | **p** |
| **day 1-3** | 0.177 | [0, 0.452] | 1.47E-06 | 0.321 | [0.003, 0.651] | 4.86E-13 |
| **day 2-4** | 0.276 | [0, 0.612] | 3.57E-10 | 0.401 | [0.005, 0.714] | 8.06E-17 |
| **day 3-5** | 0.229 | [0, 0.51] | 3.62E-08 | 0.363 | [0, 0.702] | 1.74E-14 |
| **day 4-6** | 0.182 | [0, 0.491] | 1.68E-05 | 0.297 | [0, 0.604] | 3.87E-10 |
| **day 5-7** | 0.19 | [0, 0.459] | 2.15E-05 | 0.307 | [0, 0.663] | 1.37E-10 |
